# A programmable mRNA platform for miRNA detection via miRNA-mRNA_2_ triplex-mediated ribosomal frameshifting

**DOI:** 10.1101/2025.11.24.690164

**Authors:** Yanyu Chen, Wanqi Zhao, Huizhen Chen, Hanting Zhou, Shijian Fan, Hongyu Duan, Yuze Dai, Rongguang Lu, Chenzhong Li, Cheng Jiang, Ho Yin Edwin Chan, Gang Chen

## Abstract

Programmed −1 ribosomal frameshifting (−1 PRF) is a recoding mechanism utilized by viruses to expand their coding capacity and modulate the stoichiometric ratio of −1 frame and 0 frame translation products. The stability of mRNA secondary structure at the ribosomal entry site within the frameshifting stimulating elements (FSEs) determines the frameshifting efficiency. Here, we report the development of a programmable mRNA-based platform that detects specific mature microRNA (miRNA or miR) by converting their presence into a quantifiable protein output through miRNA-triggered −1 PRF. We designed a triplex-forming mRNA (TF-mRNA) platform to selectively trap target miRNAs through the formation of major-groove mRNA-miRNA-mRNA (miR-mRNA_2_) triplexes. Bio-layer interferometry and fluorescence binding studies confirmed that TF-mRNA forms stable complexes with cognate miRNAs with low nanomolar affinity and prolonged dissociation rate. Critically, the formation of miR-mRNA_2_ triplex robustly stimulated ribosomal frameshifting in a cell-free dual-luciferase translation system, acting as a miRNA-dependent molecular switch. The generality of this TF-mRNA platform has been verified for several disease-associated purine-rich miRNAs, and it is suitable for targeting a wide range of other purine-enriched miRNAs. This programmable TF-mRNA platform establishes a foundation for developing novel diagnostic tools and synthetic biology circuits that convert the presence of miRNA into a quantifiable protein output.

**TOC Figure:** 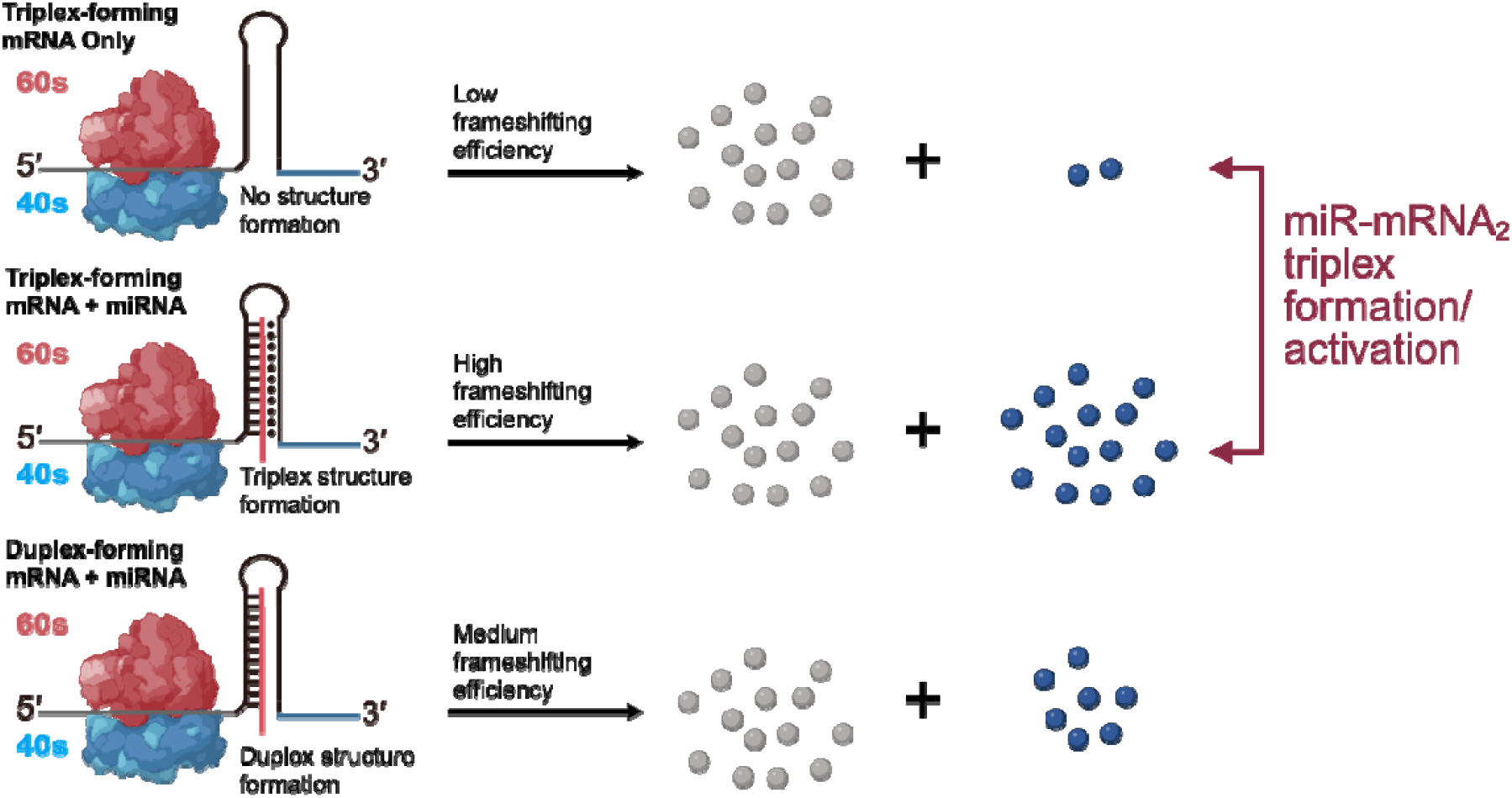

## INTRODUCTION

Programmed −1 ribosomal frameshifting (−1 PRF) is an important mechanism for viruses’ gene expression regulation **(Figure 1a)** *(1)*. During −1 PRF, ribosomes move one nucleotide backward, from 0 frame to −1 frame along the coding sequence on the mRNA, ultimately resulting in changes in protein expression **(Figure 1a, b)**. It has been reported that the stability of mRNA secondary structures (such as hairpins, pseudoknots, G-quadruplexes, etc.) downstream of the slippery sequence determines the frameshifting efficiency *(2-9)*. It was also suggested that the conformational dynamics of mRNA structures at the entry site contribute to ribosomal frameshifting *(10, 11)*.

**Figure 1.**
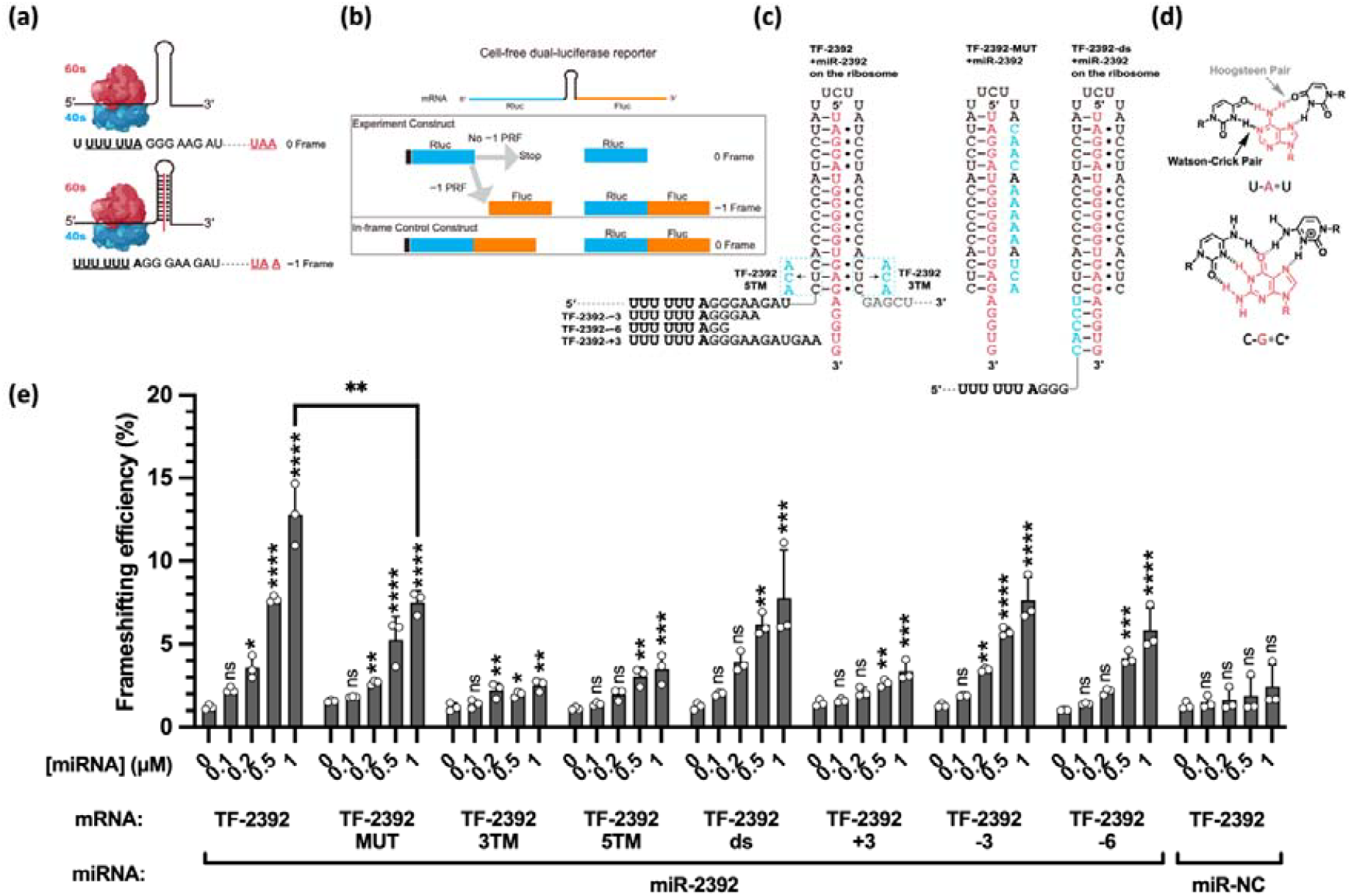
The intermolecular triplex formed by miR-2392 and TF-2392 can strongly stimulate programmed −1 ribosomal frameshifting. **(a)** Schematic diagrams of programmed −1 ribosomal frameshifting. **(b)** Workflow of cell-free dual-luciferase assay. **(c)** Schematic diagrams of TF-2392 and its mutant on the ribosome. **(d)** Chemical structures of C-G•C^+^ and U-A•U triples. **(e)** Frameshifting efficiency stimulated by miR-mRNA_2_ triplex formation. The concentration of mRNA is 20 nM. The data were analyzed by GraphPad Prism 10 and calculated by an ordinary one-way analysis of variance (ANOVA) using Dunnett’s multiple comparisons test against the mean of the mRNA alone group. The error bars represent ± S.D. * P<0.05, ** P<0.01, *** P<0.001, **** P<0.0001, ns: not significant.

An RNA triplex contains a Watson-Crick duplex bound to a third strand *(12, 13)*. Typically, a major-groove RNA triplex comprises pyrimidine•purine-pyrimidine base triples (“-”: Watson-Crick and “•” Hoogsteen base pair) such as U•A-U and C^+^•G-C base triples **(Figure 1a, c, d)** *(13)*. Intramolecular RNA major-groove and minor-groove triplex formation within a pseudoknot (RNA tertiary interactions) can significantly stimulate ribosomal frameshifting, a recoding mechanism many viruses utilize to express their replicase *(5-7, 14)*. We previously demonstrated that an intermolecular major-groove triplex formed between double-stranded RNA (dsRNA) and dsRNA-binding peptide nucleic acid (dbPNA) stimulates ribosomal frameshifting by enhancing the stability of mRNA duplex structure at the ribosomal entry site *(15, 16)*.

Reporter gene systems that enable the translation of fluorescent or luminescent proteins are useful for studying gene regulation mechanisms, including ribosomal frameshifting **(Figure 1b)**, as well as biosensing and synthetic biology applications *(17-19)*. We hypothesize that intermolecular RNA duplex and triplex formation may facilitate enhanced ribosomal frameshifting, which may provide the foundation for studying and detecting biologically important RNAs through the readout signals of proteins through ribosomal frameshifting.

The microRNAs (miRNAs or miRs) are small non-coding RNAs (∼22 nucleotides in length, see **Figure 1c** for example) that modulate gene expression by guiding the argonaute (AGO) in targeting the complementary sequences of 3′ untranslated regions (UTR) of mRNAs, resulting in mRNA translational inhibition and degradation *(20-23)*. Various miRNAs have been demonstrated to be dysregulated in cancer, infectious diseases, and other diseases *(24-26)*. Therefore, miRNAs and their hairpin precursors become promising therapeutics and diagnostic markers that are targeted by oligonucleotides, such as locked nucleic acids (LNAs), peptide nucleic acid (PNA), and other chemically modified nucleic acid analogs *(24, 27-33)*. However, the potential of using an intermolecular RNA triplex, formed between a synthetic mRNA and an endogenous miRNA, to stimulate PRF for detection purposes remains unexplored. We hypothesized that designing triplex-forming mRNAs (TF-mRNAs) could allow selectively trapping of specific miRNAs and converting their presence into a measurable frameshifting signal **(Figure 1a-c)**. In this study, we have revealed that TF-mRNA can indeed selectively form stable triplexes with free miRNAs using the bio-layer interferometry (BLI) and fluorescence binding methods. Furthermore, the mRNA-miRNA-mRNA (miR-mRNA_2_) triplex significantly stimulates ribosomal frameshifting efficiency in cell-free dual-luciferase translation systems. The work builds the foundation for developing a miRNA detection system and miRNA-based gene regulatory system in synthetic biology.

## RESULTS AND DISCUSSION

### TF-2392 forming a triplex with miR-2392

It was previously reported that miR-2392 is upregulated and suppresses mitochondrial activity during SARS-CoV-2 infection progression *(26)*. Thus, miR-2392 may serve as a diagnostic biomarker and antiviral drug target *(26)*. The mature miR-2392 contains an abundance of purine of 80% (purine number/total base number), which is an ideal target for forming a miR-mRNA_2_ triplex **(Figure 1c)** *(13)*. We designed a TF-mRNA forming a parallel major-groove triplex with miR-2392 **(Figure 1c)**. In addition, an mRNA mutant with the mutagenesis of the 3′ arm sequences of TF-2392 (TF-2392-MUT) was also designed, which is expected to completely disrupt the triplex formation by the disruption of the Hoogsteen interactions **(Figure 1c)**.

BLI was employed to determine the binding parameters, including dissociation constant (*K*_D_), association rate constant (*k*_on_), and dissociation rate constant (*k*_off_), using biotin-labelled short RNA constructs **(Figure 2a**,**b)** *(15)*. The BLI assay was performed in 200 mM NaCl, 0.5 mM EDTA, with or without 5 mM MgCl_2_, and 20 mM MES (pH 5.5) or HEPES (pH 7.5/8.5), respectively. Consistent with previous research on RNA•RNA_2_ triplexes *(34-40)*, the binding affinity was decreased with increasing pH; the *K*_D_ values are 0.91±0.02, 1.30±0.02, and 2.48±0.03 nM, at pH 5.5, 7.5, and 8.5, respectively (**Figure 2b,d, S1**), as lowering the pH promotes protonation of the cytidine N3 nitrogen, thereby facilitating C^+^•G-C base triple formation **(Figure 1d)** *(41)*. The BLI data show that the increase in *K*_D_ (lower affinity) with higher pH was primarily driven by an increase in the dissociation rate (*k*_off_) while the association rate (*k*_on_) remained relatively unchanged **(Figure 2d, S1)**. The addition of Mg^2+^ resulted in a decrease in the *K*_D_ value, consistent with the reported work that Mg^2+^ stabilized the triplex structure *(13)* (**Figure 2b,d**). Moreover, compared to TF-2392, TF-2392-MUT exhibits a significant increase in the *K*_D_ and *k*_off_ values, particularly at pH 5.5, with a ∼ten-fold change observed (**Figure 2d**).

**Figure 2.**
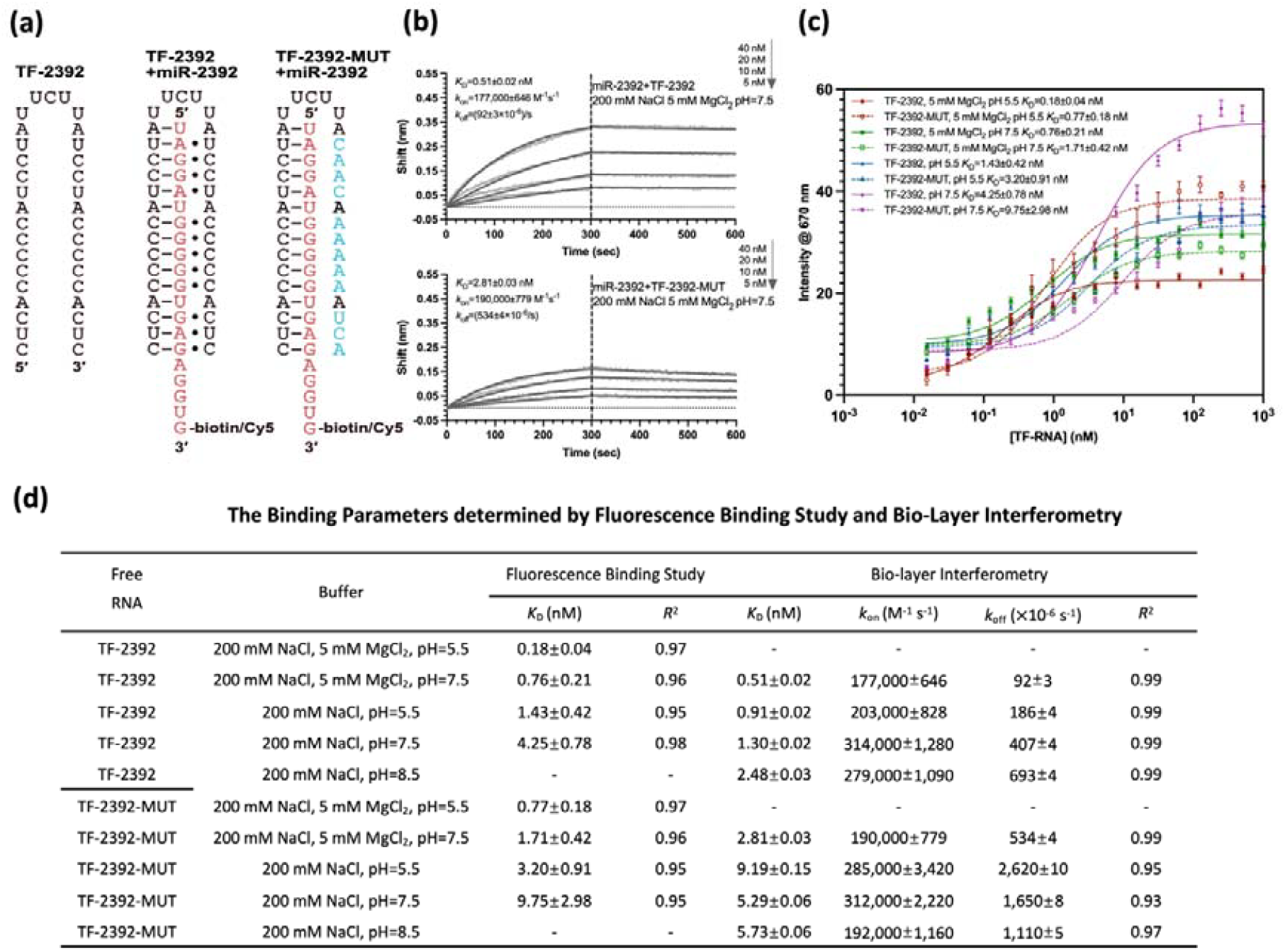
miR-2392 binds to TF-2392 mRNA through triplex formation. **(a)** Schematic diagrams of TF-2392 and its mutant with miR-2392. **(b)** Representative binding data of miR-2392 with TF-2392 and TF-2392-MUT based on bio-layer interferometry assay. **(c)** Representative binding data of miR-2392 with TF-2392 and TF-2392-MUT based on fluorescence binding study. **(d)** The summary of binding parameters obtained by BLI and fluorescence binding study, in the buffer, 200 mM NaCl, 0.5 mM EDTA, with or without 5 mM MgCl_2_, and 20 mM MES (pH 5.5) or HEPES (pH 7.5/8.5). The error bars for BLI data represent ± S.E.M for the representative data. The error bars for the fluorescence binding data represent ± S.D.

To further confirm the binding parameters obtained by BLI. The fluorescence binding study method was utilized (with the replacement of the biotin label with a Cy5 dye, **Figure 2a**) to verify the binding in parallel. The fluorescence binding assay was conducted in 200 mM NaCl, 0.5 mM EDTA, with or without 5 mM MgCl_2_, and 20 mM MES (pH 5.5) or HEPES (pH 7.5), respectively. The addition of TF-2392/TF-2392-MUT results in a dose-dependent increase of Cy5 fluorescence intensity, which facilitates the quantification of the binding affinities in solution (**Figure 2c, d, S2**). The trends of *K*_D_ values are consistent with BLI results (**Figure 2c, d, S2**). For TF-2392, the *K*_D_ values show a pH-dependent manner, and the presence of Mg^2+^ enhances the binding affinity. TF-2392-MUT exhibits higher *K*_D_ values in all conditions compared to TF-2392. To sum up, the data establish that the 3′ arm Hoogsteen pair-forming sequence of TF-2392 is crucial for the stable complex structure formation between TF-2392 and miR-2392.

To directly probe the Hoogsteen pair formation, we designed an RNA hairpin (HP-2392) that covalently links miR-2392 and the 5′ arm of TF-2392, with the 3′ arm of TF-2392 as a triplex-forming oligonucleotide (TFO, designated as TFO-WT) potentially binding to HP-2392 through Hoogsteen base pairing only **(Figure 3a, Table S1)**. BLI was also employed to determine the binding parameters. The experiments were conducted at 200 mM NaCl at pH 5.5, with or without 5 mM MgCl_2_. The triplex formations between HP-2392 and TFO-WT can be detected with *K*_D_ values of 245±3 nM (with MgCl_2_) and 504±7 nM (without MgCl_2_), respectively **(Figure 3b, d, S3)**. As expected, a significantly weakened binding was observed for HP-2392 and TFO-MUT **(Figure 3c, d, S3)**.

**Figure 3.**
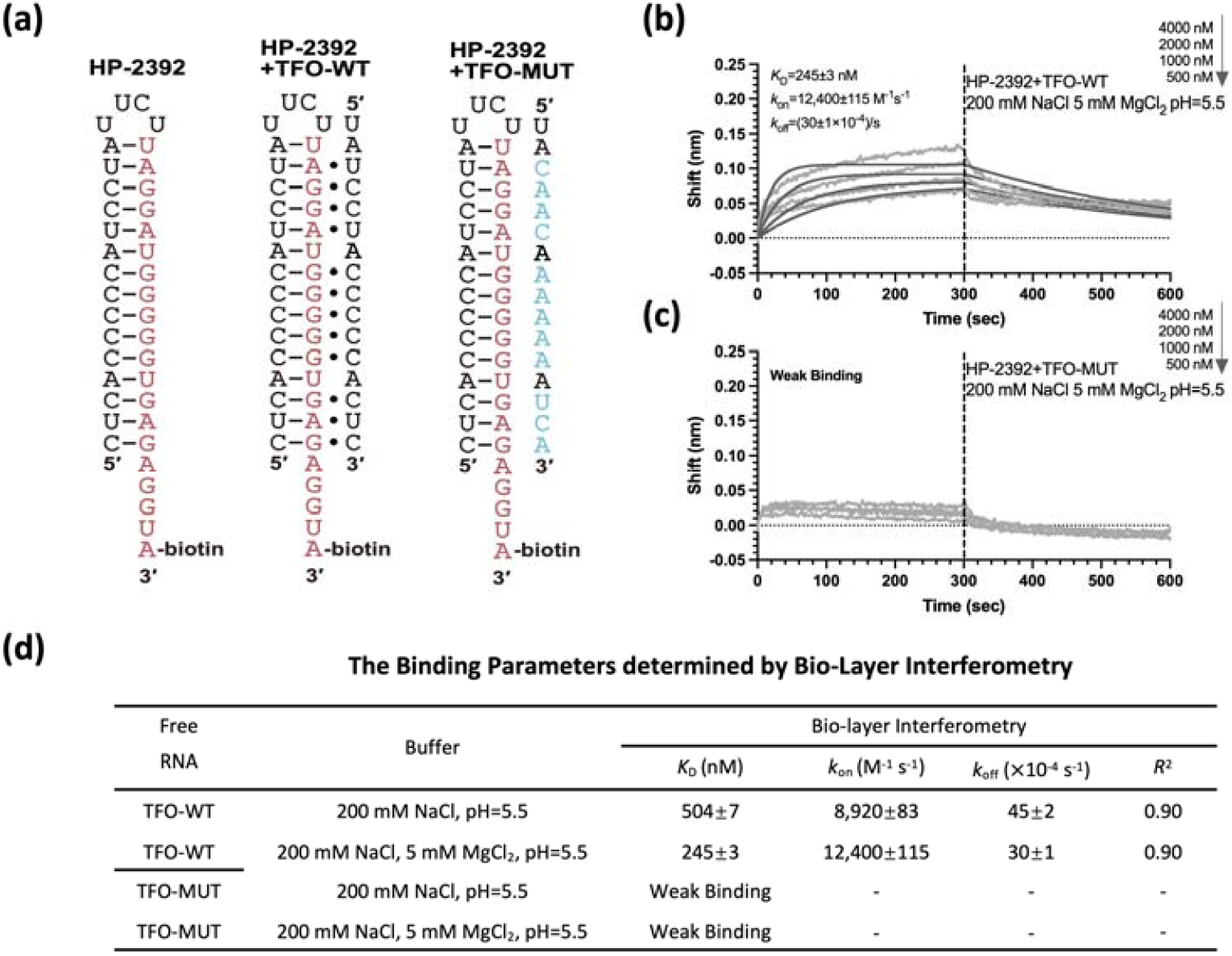
The triplex formation can be further proved by the binding data of HP-2392 with TFO-WT and TFO-MUT. **(a)** Schematic diagrams of HP-2392 binding with TFO-WT and its mutant. **(b**,**c)** Representative binding data of HP-2392 binding with TFO-WT and TFO-MUT, based on BLI. **(d)** The summary of binding parameters obtained by BLI, in the buffer, 200 mM NaCl, 0.5 mM EDTA, with or without 5 mM MgCl_2_, and 20 mM MES at pH 5.5. The error bars for BLI data represent ± S.E.M for the representative data.

We further probed triplex formation by assessing structural invasion *(7, 42, 43)*. We designed a DNA sequence that is expected to invade the TF-miR complexes **(Figure S4)**. The BLI results showed that, compared with the mutant complex (TF-2392-MUT with miR-2392, with a *K*_D_ value of 194±3 nM) with a weakened Hoogsteen base pairing interaction, the TF-2392/miR-2392 complex was harder to invade, with a larger *K*_D_ value (250±4 nM) **(Figure S4)**. Taken together, our data suggest that the designed triplex-forming mRNA binds to miR-2392 tightly through both Watson-Crick and Hoogsteen pairing, which may stabilize the structure and enhance its ribosomal frameshifting efficiency.

### miR-mRNA_2_ triplex formation stimulates frameshifting in a cell-free system

To determine the frameshift-stimulating activity of TF-2392, the dual-luciferase reporter system **(Figure 1b)** was employed based on the p2luc plasmid *(15, 16, 44)*. The sequences in **Figure 1c** were inserted into p2luc following an upstream slippery site (UUUUUUA) and an eight-nucleotide spacer (GGGAAGAU) **(Figure 1c, S5)**. The schematic of the dual-luciferase reporter is shown in **Figure 1b**, and the *Renilla* luciferase and *Firefly* luciferase are 0-frame and −1-frame translation products, respectively.

Herein, an expected triplex, as shown in **Figure 1a-c**, is hypothesized to stimulate ribosomal frameshifting by temporarily inhibiting the ribosome helicase activity *(45, 46)*. Thus, −1 PRF, in this study, is expected to serve as a tool for the detection of the triplex formation. We observed that, in the absence of miR-2392, the frameshifting efficiencies of TF-2392 and TF-2392-MUT are 1.2±0.1% and 1.6±0.1%, respectively **(Figure 1e)**. Upon the application of free miR-2392, we observed a miR-2392 concentration-dependent activity in stimulating frameshifting efficiency **(Figure 1e)**. The TF-2392 showed the highest frameshifting efficiency (12.81±1.86%) with 1000 nM miR-2392. However, the efficiency was decreased to 7.51±0.70% for TF-2392-MUT with 1000 nM miR-2392, indicating that the 3′ arm of TF-2392 plays a critical role in forming the miR-mRNA_2_ triplex during the translation recoding process. Moreover, the frameshifting efficiency and normalized expression level of Fluc are positively correlated with *K*_D_ and *k*_off_ values **(Figure S6)**. It was previously reported that an antisense LNA can stimulate frameshifting efficiency with a length of ∼12-18 bases *(47)*, consistent with our data showing that the miR-mRNA duplex can also stimulate frameshifting efficiency, although with a lower efficiency than that of the miR-mRNA_2_ triplex. Taken together, our data clearly show that, compared to intermolecular duplex formation, the formation of an intermolecular triplex structure significantly stimulates ribosomal frameshifting.

### The spacer length between the slippery sequence and intermolecular triplex affects frameshifting

Next, we determined the influence of the length of the single-stranded spacer sequence on frameshifting stimulated by miR-mRNA_2_ triplex formation. We designed a series of TF-mRNA constructs with eleven, five, and two-nucleotide spacer sequences (TF-2392+3, TF-2392−3, and TF-2392−6) as shown in **Figure 1c**. Dual-luciferase reporter assay data illustrate that the baseline frameshifting efficiencies of TF-2392+3, TF-2392−3, and TF-2392−6 are 1.45±0.19%, 1.31±0.11% and 1.03±0.02%, respectively. We observed a miR-2392 concentration-dependent activity in stimulating frameshifting efficiency, but generally weaker than that of TF-2392 **(Figure 1e)**. The efficiency was decreased to 3.38±0.60%, 7.64±1.37% and 5.86±1.33% for TF-2392+3, TF-2392−3, and TF-2392−6 with 1000 nM miR-2392, respectively. Compared with the eight-nucleotide spacer, constructs with a shortened or lengthened spacer exhibited a reduced frameshifting stimulation. Interestingly, extending the spacer sequences lowers the frameshifting efficiency more significantly than shortening them **(Figure 1e)**, which agrees with a previous report *(48)*.

Mutating the terminal three bases in both the 5′ arm (TF-2392-5TM) and 3′ arm (TF-2392-3TM) results in a significant reduction in frameshifting efficiency **(Figure 1e)**, which suggests that both the terminal Watson-Crick and Hoogsteen base pairs (at the entry of the ribosome) play a critical role in stimulating ribosomal frameshifting. Surprisingly, extending the Watson-Crick duplex by base pairing with the complete miR-2392 sequence (TF-2392-ds) results in a frameshifting efficiency in between TF-2392 and TF-2392−3, suggesting that positioning triplex at the ribosome entry site is critical in stimulating high frameshifting efficiency. It has been reported that Watson-Crick duplex can stimulate ribosomal frameshifting, ideally with a shortened single-stranded spacer with about 2-3 bases *(46)*. We speculate that for TF-2392-ds, although the overall downstream structural stability is enhanced with the intermolecular triplex intact, extending the Watson-Crick duplex causes the shortening of the single-stranded spacer to three bases, resulting in the frameshifting stimulation mainly determined by the Watson-Crick duplex region at the ribosome entry site, instead of the triplex formed 5-base pair further downstream. This finding underscores that optimal frameshifting stimulation requires precise positioning of the triplex structure at the ribosomal entry site, rather than merely enhancing the overall downstream mRNA structure stability.

### The general applicability of the TF-mRNA platform to target other miRNAs

We subsequently evaluated the generality of the TF-mRNA platform on other purine-enriched miRNAs. We chose several miRNAs with purine abundance higher than 60%. miR-142 (62%), miR-198 (77%) and miR-6124 (100%) were selected. These miRNAs have been reported to be associated with the progression of some cancers *(49, 50)* and the wound healing process *(51)*. Three mRNAs, TF-142, TF-198, and TF-6124, were designed in a manner similar to TF-2392 **(Figure 4a)**. The baseline frameshifting efficiency values of TF-142, TF-198 and TF-6124 are 2.20±0.05%, 1.32±0.08% and 1.18±0.16%, respectively. The frameshifting assay data also showed miRNA concentration-dependent activities in stimulating frameshifting efficiency **(Figure 4b-d)**. By adding 2000 nM of the miRNAs, the frameshifting efficiencies rose to 3∼8-fold higher than their respective baseline efficiency values. For example, TF-198 shows an 8-fold stimulation of the frameshifting efficiency with 8.89±0.60% by adding 2000 nM miR-198. The normalized FLuc levels also rose in a concentration-dependent manner **(Figure S7)**.

**Figure 4.**
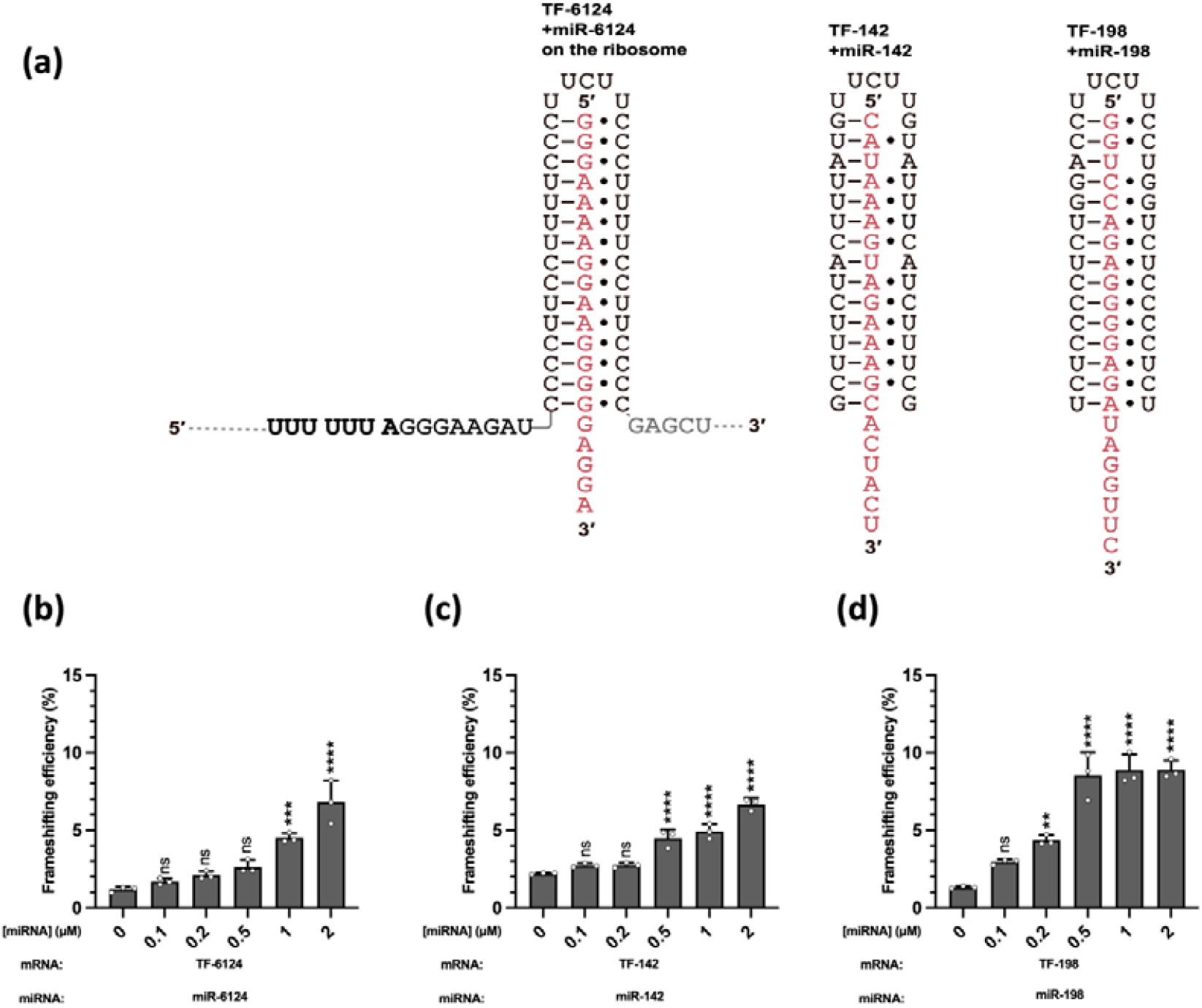
The general applicability of the TF-mRNA platform to target other miRNAs. **(a)** Schematic diagrams of TF-6124, TF-142 and TF-198 on the ribosome. **(b-d)** Frameshifting efficiency stimulated by miR-mRNA_2_ triplex formation. The mRNA concentration is 20 nM. The data were analyzed by GraphPad Prism 10 and calculated by an ordinary one-way analysis of variance (ANOVA) using Dunnett’s multiple comparisons test against the mean of the mRNA alone group. The error bars represent ± S.D. * P<0.05, ** P<0.01, *** P<0.001, **** P<0.0001, ns: not significant.

We speculate that the reduced frameshifting efficiency values compared to those of TF-2392 may be due to the varied stability of the terminal base triples, as observed in duplex-stimulated frameshifting *(4)*. In TF-2392, the three terminal base triples include two C^+^•G-C base triples flanking a U•A-U triple. However, TF-6124 contains three consecutive C^+^•G-C base triples, which may not be thermodynamically favorable for stable base triple formation with three consecutive protonated C bases. TF-142 has a terminal Watson-Crick pair without base triple formation. In TF-198, the three terminal base triples include a C^+^•G-C base triple flanked by two U•A-U triple. Clearly, two C^+^•G-C base triples flanking a U•A-U triple may be ideal for positioning the triplex at the mRNA entry site for optimal stimulation of frameshifting *(52)*.

## CONCLUSIONS

In this study, we designed a programmable TF-mRNA platform and systematically investigated the triplex formation between TF-mRNA constructs and the miRNAs, as well as the *in vitro* biological activities in a cell-free system. We demonstrate that TF-mRNA can form triplexes robustly with target miRNAs, and this triplex formation can be detected by ribosomal frameshifting stimulation. Moreover, the TF-mRNA platform shows potential applications for targeting other miRNAs, especially purine-enriched miRNAs. Importantly, a single-stranded spacer of approximately 8 bases positions the downstream intermolecular triplex at the mRNA entry site, which is ideal for frameshifting stimulation. This work highlights the potential of RNA triplexes as control elements in synthetic biology. Further work is needed to explore multiplexed detection schemes and test whether triplex formation affects AGO loading and the platform’s performance in the cellular environment. The findings from our work provide valuable references for future work on developing mRNA-based diagnosis tools, engineered genetic circuits that respond to RNA-based signals, and potential RNA-targeting therapeutics.

## MATERIALS AND METHODS

### Bio-layer interferometry

The binding kinetics of TF-2392/TF-2392-MUT to biotinylated miR-2392 were analyzed by bio-layer interferometry (BLI) using a Gator® Prime instrument (Gator Bio) at 30°C, with 300-second association and dissociation phases. Streptavidin (SA) biosensors were employed to immobilize miRNA diluted to 200 nM in binding buffer (200 mM NaCl, 0.5 mM EDTA with or without 5 mM MgCl_2_, 20 mM MES (pH 5.5) or HEPES (pH 7.5/8.5). TF-2392/TF-2392-MUT samples were serially diluted to 40, 20, 10, and 5 nM in the same buffer. For assays testing the binding affinity of HP-2392 and TFO-WT/TFO-MUT, the concentration of HP-2392 was 200 nM, while the concentration of TFO-RNAs was adjusted to 4000, 2000, 1000, and 500 nM. The HP-2392 RNA was annealed by heating to 95°C for 5 min followed by snap-cooling on ice. For the invasion assay, the immobilized miRNAs were first bound with 40 nM TF RNAs in the binding buffer, then invaded by the INV DNA **(Table S1)** with 4000 and 2000 nM. Binding parameters (*K*_D_, *k*_on_ and *k*_off_) were derived from both Global and unlinked fitting models. Real-time binding curves (Δλ shift vs. time, 0.1-s resolution) and fitted plots were processed using GraphPad Prism 10 to generate the figures.

### Fluorescence Binding Study

The fluorescence binding assay was employed to analyze the binding affinity of TF-2392/TF-2392-MUT with miR-2392. The method is similar to what we described previously *(53)*. miR-2392-Cy5, TF-2392, and TF-2392-MUT were dissolved in 200 mM NaCl, 0.5 mM EDTA with or without 5 mM MgCl_2_, 20 mM MES (pH 5.5) or HEPES (pH 7.5) prior to use. The RNA oligonucleotides were annealed at 95□°C for 5□min before the experiment. The miR-2392-Cy5 (0.01□nM) was then mixed with varying concentrations of RNA in different binding buffers. The mixtures were incubated at 25□°C in the dark for 30 min. Fluorescence intensity scans were performed using a Spark® multimode microplate reader with an excitation wavelength of 620 nm (bandwidth, 10□nm) and emission detection from 650□nm to 750□nm. Parallel (Iperpendicular) and perpendicular (Iperpendicular) fluorescence intensities were recorded using SparkControl software (version 2.1). Binding affinities (*K*_D_) were calculated by non-linear regression using OriginPro 2025, as previously described *(53, 54)*. The figures were generated using GraphPad Prism 10.

### Plasmid construction

The frameshifting efficiency was quantified using the p2luc dual-luciferase reporter plasmid, generously provided by Prof. Samuel E. Butcher *(4)*, wherein the TF-RNAs’ DNA sequences were inserted between the *Renilla* and *firefly* luciferase coding regions via BamHI and SacI restriction sites, as previously established *(15, 16)*. A modified in-frame control (IFC) plasmid was generated by disrupting the slippery sequence to enforce 100% translational fidelity *(16)*. All plasmids were purified using the Monarch Plasmid Miniprep Kit (NEB), followed by ethanol precipitation, with DNA quality and concentrations verified by a NanoDrop One spectrophotometer (Thermo Scientific).

### DNA linearization, transcription and RNA purification

Linearized DNA was generated via polymerase chain reaction (PCR) using primers p2luc-631 and p2luc3431 *(15)* and the KOD-Plus-Neo kit (TOYOBO). Each 50 µL reaction contained 5 µL 10× KOD-Plus-Neo Buffer, 5 µL 2 mM dNTPs, 3 µL 25 mM MgSO□, 1.5 µL each of 10 pmol/µL forward and reverse primers, 2.5 µL 10 ng/µL DNA template, 1 µL 1.0 U/µL KOD-Plus-Neo polymerase, and 30.5 µL nuclease-free H□O (Sangon). Reactions were mixed thoroughly and cycled in a BioRad thermocycler under the following conditions: 95°C for 30 s; 35 cycles of 95°C for 30 s, 57°C for 30 s, and 72°C for 2 min; followed by a final 72°C elongation for 2 min. PCR products were ethanol-precipitated, quantified via NanoDrop (Thermo Scientific), and transcribed using the HiScribe T7 High-Yield RNA Synthesis Kit (NEB). After 2 h of transcription at 37°C, DNA templates were digested with DNase I (NEB), and RNA transcripts were purified using an RNA cleanup kit (NEB) and resuspended in nuclease-free H□O for downstream applications.

### *In vitro* translation, dual-luciferase reporter assay and data analysis

The cell-free translation system was adapted from established methods *(15, 16)* with minor modifications. Briefly, mRNA transcripts (final concentration: 20 nM) were pre-mixed with serial dilutions of miRNAs (final concentration: 0, 0.1, 0.2, 0.5, 1 μM) in DEPC-treated water (Sangon). Each 12.5 μL reaction consisted of 8.75 μL nuclease-treated rabbit reticulocyte lysate (RRL, Promega), 1.25 μL miRNA solution, and 2.5 μL mRNA, incubated at 30°C for 90 min in a thermal cycler. For luminescence quantification, 2 μL of translation products were combined with 50 μL Dual-Glo *Firefly* Luciferase Substrate (Promega), and relative luminescence units (RLU) were immediately measured using an EnVision Multimode Plate Reader (PerkinElmer). Subsequently, 50 μL Dual-Glo Stop & Glo *Renilla* Substrate (Promega) was added to quantify RLU for normalization. All conditions were tested in triplicate, with data analyzed in GraphPad Prism. The formula for calculating the frameshifting efficiency is *(44)*:

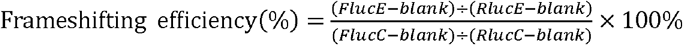

## AUTHOR INFORMATION

### Authors

Yanyu Chen – School of Medicine, The Chinese University of Hong Kong, Shenzhen (CUHK-Shenzhen), Shenzhen, 518172, Guangdong, P. R. China.

Wanqi Zhao – School of Medicine, The Chinese University of Hong Kong, Shenzhen (CUHK-Shenzhen), Shenzhen, 518172, Guangdong, P. R. China.

Huizhen Chen – School of Life Sciences, The Chinese University of Hong Kong, Shatin N.T., Hong Kong SAR, P. R. China.

Hanting Zhou – School of Medicine, The Chinese University of Hong Kong, Shenzhen (CUHK-Shenzhen), Shenzhen, 518172, Guangdong, P. R. China.

Shijian Fan – School of Medicine, The Chinese University of Hong Kong, Shenzhen (CUHK-Shenzhen), Shenzhen, 518172, Guangdong, P. R. China.

Hongyu Duan – School of Medicine, The Chinese University of Hong Kong, Shenzhen (CUHK-Shenzhen), Shenzhen, 518172, Guangdong, P. R. China.

Yuze Dai – School of Medicine, The Chinese University of Hong Kong, Shenzhen (CUHK-Shenzhen), Shenzhen, 518172, Guangdong, P. R. China.

### Author Contributions

Gang Chen, Ho Yin Edwin Chan, Cheng Jiang, Rongguang Lu, and Yanyu Chen conceived the study. Yanyu Chen, Wanqi Zhao, Huizhen Chen, Hanting Zhou, Shijian Fan, Hongyu Duan, and Yuze Dai performed the experiments. Yanyu Chen, Gang Chen, Rongguang Lu, Huizhen Chen and Wanqi Zhao wrote the manuscript. All of the authors approved the final submission of the manuscript.

### Notes

The authors declare no competing financial interest.

## ASSOCIATED CONTENT

### Supporting Information

DNA and RNA oligonucleotides used in this study, binding data of TF-2392 or TF-2392-MUT with miR-2392 in different pH conditions, fluorescence spectra at 650-750 nm of TF-2392 or TF-2392-MUT binding with miR-2392, binding data of TFO-WT or TFO-MUT with HP-2392, TF-miRNA complex structural invasion data, sequencing data for the plasmids, normalized Fluc and Rluc data for TF-2392 and its derivatives, and normalized Fluc and Rluc data for other TF-mRNA constructs.

## ACKNOWLEDGEMENTS

This work was supported by a National Natural Science Foundation of China (NSFC) project (grant 22177098 to G.C.); The Chinese University of Hong Kong, Shenzhen (CUHK-Shenzhen) University Development Fund (to G.C.); a fund from the Shenzhen-Hong Kong Cooperation Zone for Technology and Innovation (HZQB-KCZYB-2020056 to G.C.); Shenzhen Natural Science Foundation in Basic Research Fund (General Program, JCYJ20240813113616021); the Shenzhen Science and Technology Innovation Committee for the Shenzhen Key Laboratory Scheme (ZDSYS20220507161600001); Guangdong Basic Research Center of Excellence for Aggregate Science; State Key Laboratory of Agrobiotechnology ITC Fund (to H.Y.E.C.); Shenzhen Medical Research Fund (A2503086 to R.L.); the Shenzhen Clinical Research Center for Tuberculosis (20210617141509001 to R.L.); Longgang District Key Laboratory of Intelligent Bio-Optoelectronics and Brain Disease Diagnosis and Treatment (to C.L.); Shenzhen Peacock Team (KQTD20240729102029016 to C.L.); National Natural Science Foundation of China (NSFC) (grant W2531016 to C.L.).

