## Supplementary material for "A programmable mRNA platform for miRNA detection via miRNA-mRNA_2_ triplex-mediated ribosomal frameshifting": SI

**Table S1. DNA and RNA oligonucleotides used in this study.**

| Oligomer | Label | Sequences (5'-3') | Type |
|---|---|---|---|
| TF-2392 | / | CUCACCCCCAUCCUAUUCUUAUCCUACCCCCACUC | RNA |
| TF-2392-MUT | / | CUCACCCCCAUCCUAUUCUUACAACAAAAAAAUCA | RNA |
| miR-2392-biotin | 3'biotin | UAGGAUGGGGGUGAGAGGUG | RNA |
| miR-2392-Cy5 | 3'Cy5 | UAGGAUGGGGGUGAGAGGUG | RNA |
| miR-2392 | / | UAGGAUGGGGGUGAGAGGUG | RNA |
| HP-2392-biotin | 3'biotin | CUCACCCCCAUCCUAUUCUUAGGAUGGGGGUGAGAGGUG | RNA |
| TFO-WT | / | UAUCCUACCCCCACUC | RNA |
| TFO-MUT | / | UACAACAAAAAAAUCA | RNA |
| miR-142 | / | CAUAAAGUAGAAAGCACUACU | RNA |
| miR-198 | / | GGUCCAGAGGGGAGAUAGGUUC | RNA |
| miR-6124 | / | GGGAAAAGGAAGGGGGAGGA | RNA |
| miR-NC | / | GGAUGGAAGAUGUGGUGGGG | RNA |
| pDL-631 | / | CACTCCCAGTTCAATTACAG | DNA |
| pDL-3431 | / | TCTTATCATGTCTGCTCGAA | DNA |
| INV-DNA | / | CACCTCTCACCC | DNA |

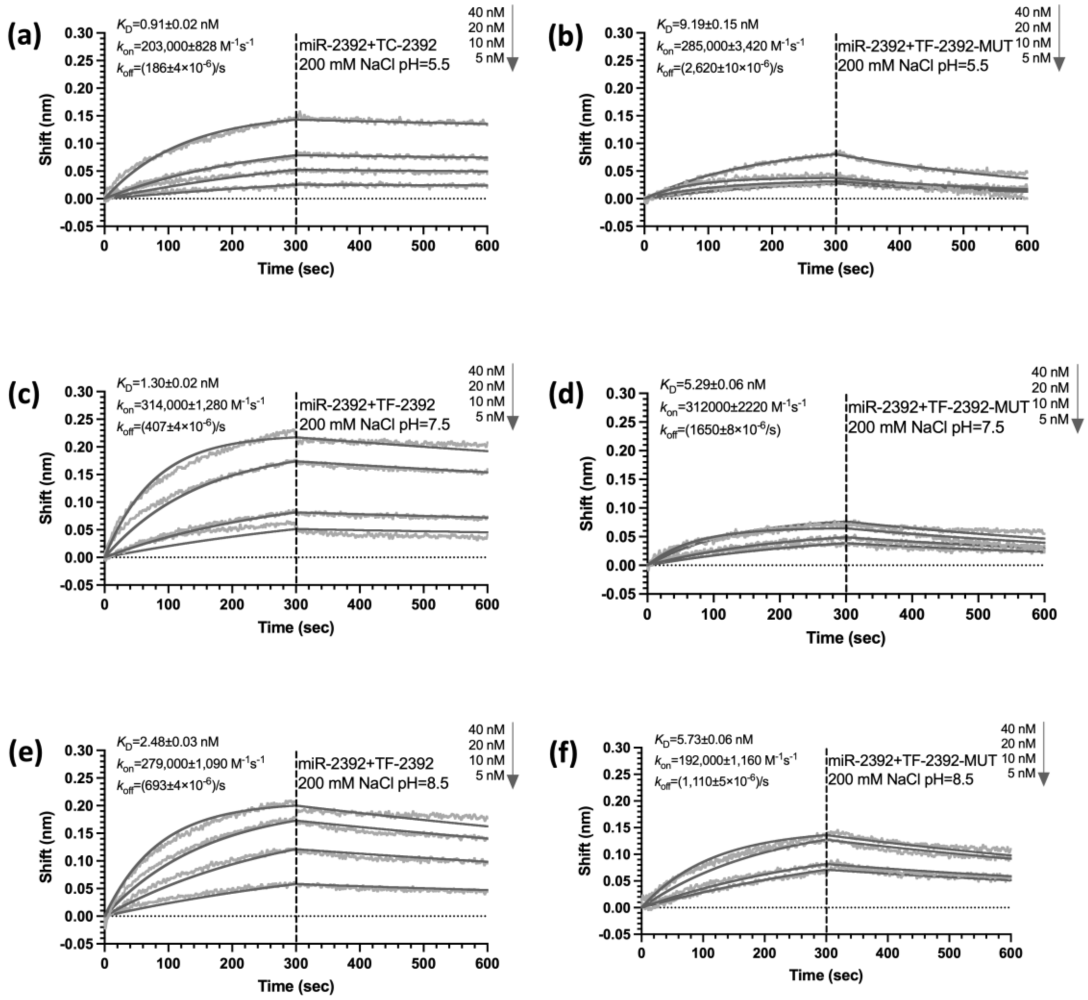


**Figure S1. Binding data of TF-2392 or TF-2392-MUT with miR-2392, obtained by BLI in different pH conditions.**

**(a,b)** pH 5.5, **(c,d)** pH 7.5, and **(e,f)** pH 8.5. Buffer containing 200 mM NaCl, 0.5 mM EDTA, and 20 mM MES (pH 5.5) or HEPES (pH 7.5/8.5). The error bars for BLI data represent ± S.E.M for the representative data.

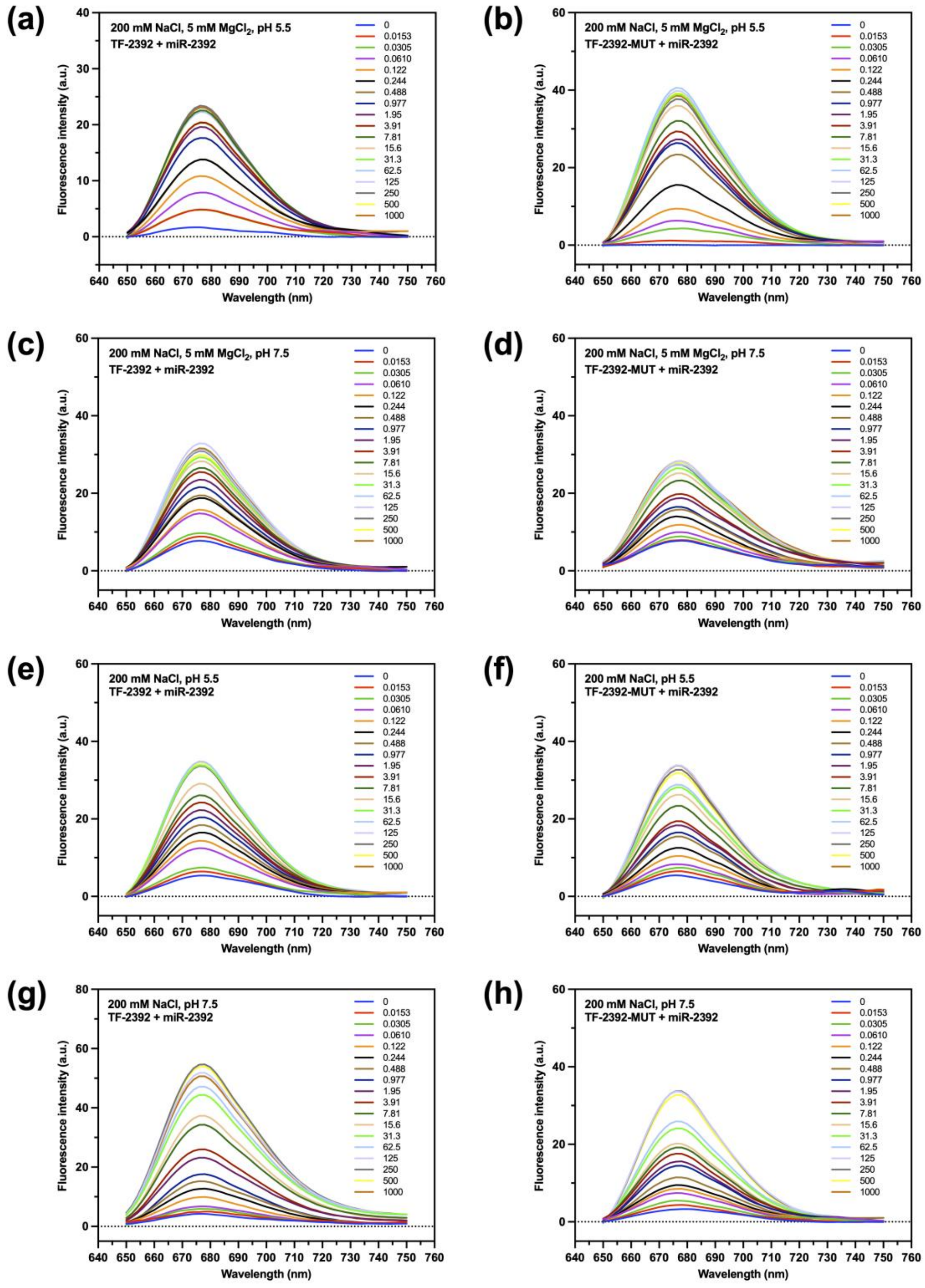


**Figure S2. Fluorescence spectra at 650-750 nm of TF-2392 or TF-2392-MUT binding with miR-2392, obtained by fluorescence binding study.**

**(a,b)** pH 5.5 with $Mg^{2+}$, **(c,d)** pH 7.5 with $Mg^{2+}$, and **(e,f)** pH 5.5 without $Mg^{2+}$ **(g,h)** pH 7.5 without $Mg^{2+}$. Buffer containing 200 mM NaCl, 0.5 mM EDTA, with or without 5 mM $MgCl_2$, and 20 mM MES (pH 5.5) or HEPES (pH 7.5)

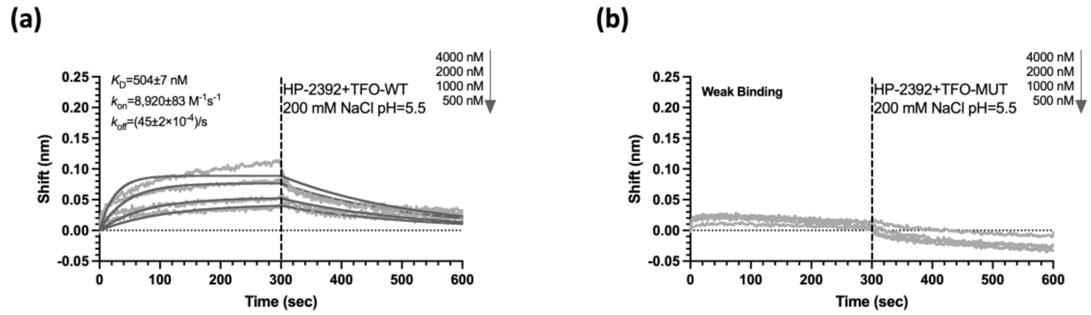


**Figure S3. Binding data of TFO-WT or TFO-MUT with HP-2392, obtained by BLI.**

**(a)** TFO-WT and HP-2392. **(b)** TFO-MUT and HP-2392. Buffer containing 200 mM NaCl, 0.5 mM EDTA, 20 mM MES at pH 5.5. The error bars for BLI data represent ± S.E.M for the representative data.

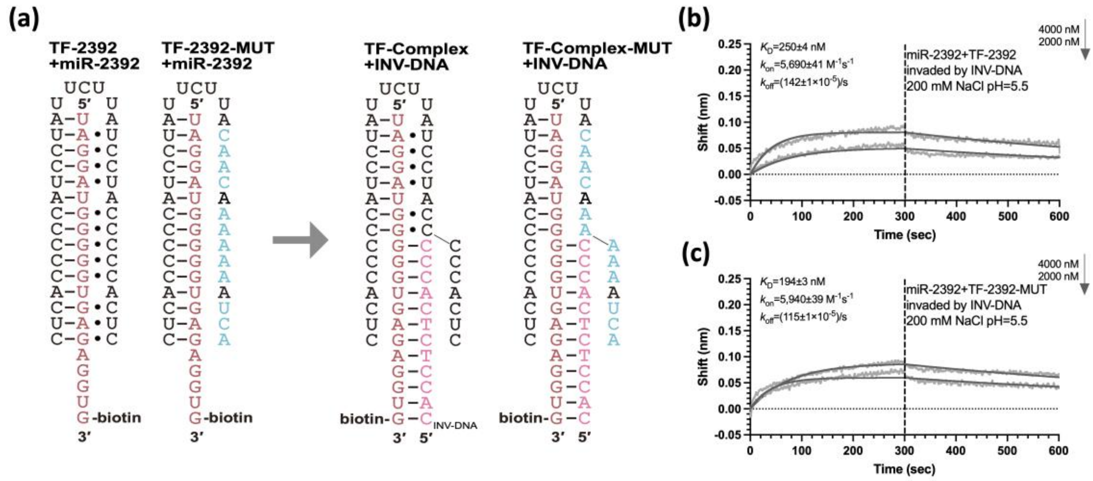


**Figure S4. The TF-2392 MUT-miR-2392 complex can be invaded by the INV-DNA more easily than the TF-2392-miR-2392 triplex.**

**(a)** Schematic diagrams of the TF-miR complexes interacting with the INV-DNA. **(b, c)** Binding data of TF-miR complexes invaded by the INV-DNA. The error bars for BLI data represent ± S.E.M for the representative data.

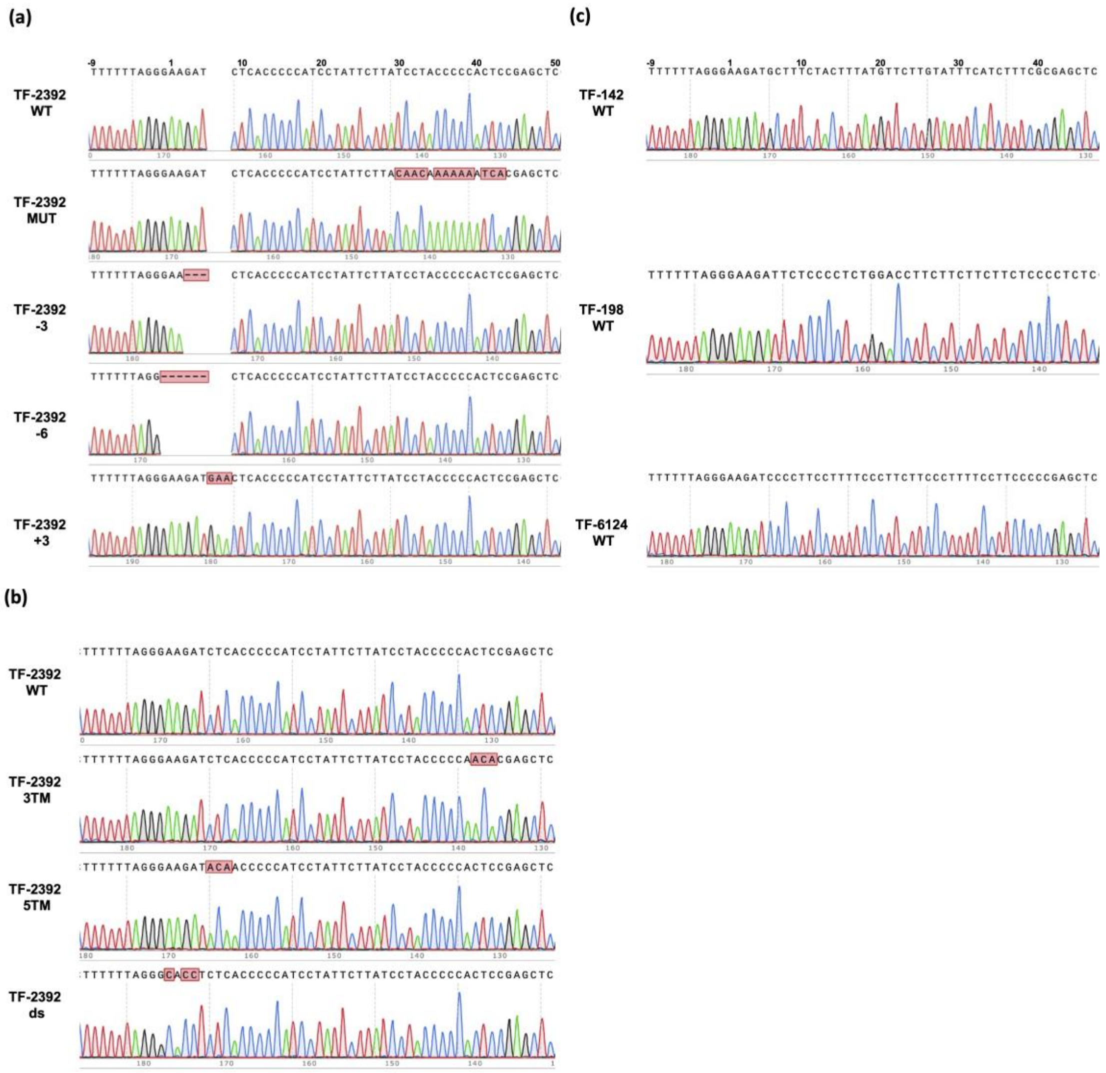

**Figure S5. Sequencing data for the plasmids.**

**(a)** Sequencing results of the plasmid TF-2392 and its derivatives, including TF-2392-MUT, TF-2392-−3, TF-2392-−6 and TF-2392-+3. **(b)** Sequencing results of the plasmid TF-2392 and its derivatives, including TF-2392-3TM, TF-2392-5TM, and TF-2392-ds. **(c)** Sequencing results of other TF-mRNA plasmids, including TF-142, TF-198, and TF-6124.

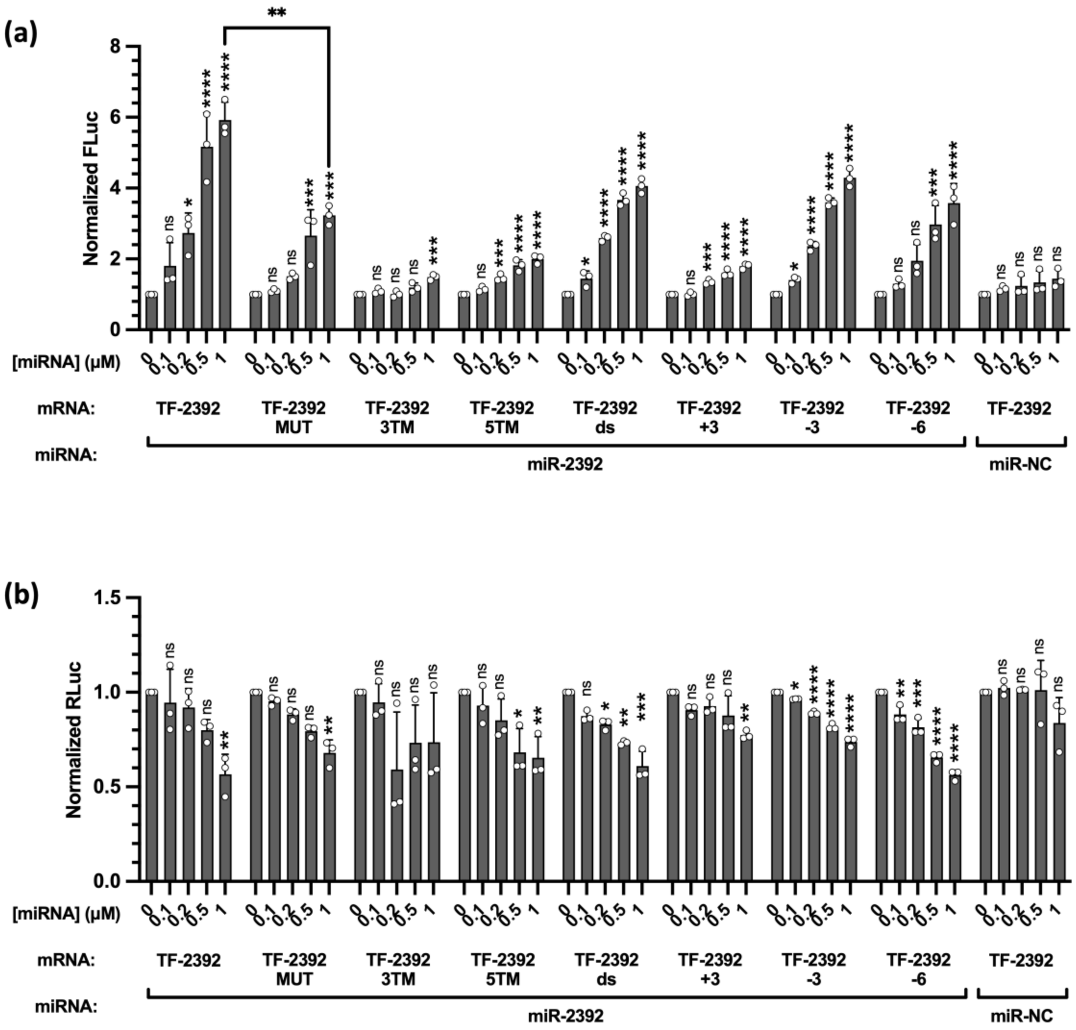


**Figure S6. Normalized Fluc and Rluc data for TF-2392 and its derivatives, obtained by cell-free dual-luciferase assay.**

**(a)** Normalised Fluc. **(b)** Normalised Rluc. The concentration of mRNA is 20 nM. The data were analyzed by GraphPad Prism 10 and calculated by an ordinary one-way analysis of variance (ANOVA) using Dunnett's multiple comparisons test against the mean of mRNA alone group. The error bars represent ± S.D. * $P < 0.05$, ** $P < 0.01$, *** $P < 0.001$, **** $P < 0.0001$, ns: not significant.

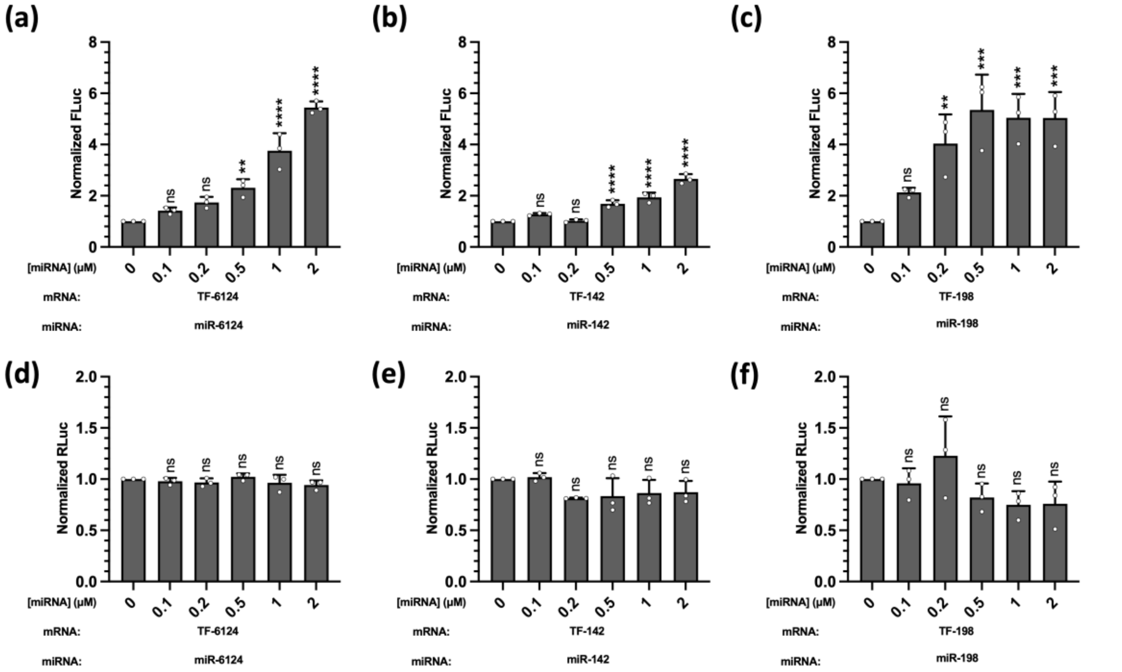


**Figure S7. Normalized Fluc and Rluc data for other TF-mRNA constructs obtained by cell-free dual-luciferase assay.**

**(a-c)** Normalised Fluc. **(d-f)** Normalised Rluc. The concentration of mRNA is 20 nM. The data were analyzed by GraphPad Prism 10 and calculated by an ordinary one-way analysis of variance (ANOVA) using Dunnett's multiple comparisons test against the mean of mRNA alone group. The error bars represent ± S.D. * $P < 0.05$, ** $P < 0.01$, *** $P < 0.001$, **** $P < 0.0001$, ns: not significant.
